# A self-consistent model for phase separation and active processes in biomolecular condensates

**DOI:** 10.64898/2026.06.01.729289

**Authors:** Marco Di Mambro, Paolo De Los Rios

**Affiliations:** Department of Biology, Institute of Biochemistry, ETH Zurich, Switzerland; Institute of Physics, École Polytechnique Fédérale de Lausanne (EPFL); Institute of Bioengineering, École Polytechnique Fédérale de Lausanne (EPFL)

## Abstract

Biomolecular condensates are thought to play a pivotal role in cellular organization by regulating biochemical reactants in space and time. Sustained molecular fluxes across condensate boundaries, together with the participation of phase-separating molecules in active chemical reactions such as ATP hydrolysis, call for a nonequilibrium description. Here, we propose a self-consistent framework in which diffusion–drift dynamics and chemical reactions are coupled through a conditional free energy, defined as the excess contribution to the chemical potential. Self-consistency is achieved by deriving this quantity from the same free-energy functional that governs molecular interactions and phase separation. We apply the framework to a minimal client–scaffold system and investigate how active chemical processes and phase separation interact at steady state. In doing so, our approach recovers the fundamental rules previously identified for the emergence of nonequilibrium steady-state fluxes. The model shows that active reactions involving the scaffold molecules can regulate the phase behavior of the condensate. Moreover, nonequilibrium steady-state fluxes are maximal near the boundary between the phase-separated and homogeneous regimes, suggesting that condensates sustaining molecular transport may operate close to their stability threshold. In the same region, client fluxes are also enhanced, revealing an indirect coupling between scaffold activity and client transport. These results provide a baseline for developing more detailed theories of chemically active condensates.

---

Biomolecular condensates allow cells to compartmentalize biochemical processes without membranes. These condensates arise from the phase separation of proteins, nucleic acids, or both, and include systems as diverse as P-bodies, stress granules, and the nucleolus^1–5^. A central function of these condensates is to regulate client molecules by controlling their local concentration and chemical environment. By selectively partitioning reactants, enzymes, or cofactors, condensates can enhance or modulate biochemical reactions through local enrichment^6–10^.

Beyond static partitioning, condensates may also contribute to directional molecular transport. Sustained fluxes of material between compartments are essential for cellular organization, and condensates have been proposed to facilitate such fluxes by coupling selective partitioning, chemical reactions, and molecular exchange across phase boundaries. This possibility appears most clearly in the nucleolus, where ribosomal components are progressively processed through distinct subcompartments during ribosome biogenesis^5,11^. More generally, the question of whether condensates regulate not only where molecules accumulate but also how they move and are chemically processed — by sustaining directed fluxes across their phase boundaries — motivates a nonequilibrium description of condensate function^12^. Chemical activity, however, is not restricted to client molecules. In many biological condensates, the scaffold components themselves undergo active reactions that are directly coupled to condensate dynamics. In the nucleolus, pre-rRNA transcription by RNA polymerase I contributes to nucleolar maintenance, and transcriptional inhibition reorganizes the nucleolus into cap-like structures^13^. Another example is provided by DEAD-box ATPases, ATP-dependent RNA chaperones that are integral to ribonucleoprotein condensates such as P-bodies and stress granules^14,15^. These proteins promote condensation in their ATP- and RNA-bound states, while ATP hydrolysis drives RNA release, turnover, and condensate remodeling^14, 16, 17^. Other condensates are remodeled by molecular chaperones, such as Hsp70 and Hsp104, which can actively (*i.e*. through a functional cycle driven by ATP hydrolysis) regulate the size, structure and material state of some condensates^18–20^ . The same molecular components thus both define the phase-separated scaffold and undergo active chemical cycles that govern its assembly and dissolution.

Chemically active condensates therefore operate on two coupled levels: the scaffold components undergo active chemical cycles that remodel the condensate itself, while the condensate in turn controls the localization and reactions of client molecules. Describing these intertwined processes requires a framework that treats phase separation, reaction kinetics, and diffusive transport within a single, consistent formalism.

Standard theoretical models of biomolecular condensates are grounded in equilibrium thermodynamics, typically through Flory–Huggins or Cahn–Hilliard-type free-energy descriptions^21–24^. Nonequilibrium effects are commonly incorporated through additional reaction terms, active turnover, or phenomenological modifications of the free-energy landscape^24–26^. While these approaches have yielded important insights, they generally lack a microscopic derivation that consistently connects diffusion, interaction-driven drift, and chemical transitions within the same framework. We recently introduced a Fokker–Planck/master-equation framework for an N-particle system evolving simultaneously in physical and chemical-state space^27^. This formulation treats diffusion, interaction-driven drift, and transitions between internal chemical states on an equal footing. Under quasi-equilibrium conditions, the diffusion–drift dynamics reduce to a Cahn–Hilliard-like free-energy description, while the full framework reveals nontrivial couplings between chemical reactions and diffusion that can sustain steady-state fluxes across condensate boundaries. Two main results emerged from that work: *i*) spatial fluxes are possible only in non-equilibrium conditions, if the chemical reaction network contains cycles and if the barriers of the reactions depend on space, but not all in the same way; *ii*) if such conditions are met, fluxes are nonetheless localized at the interface of the phase separated regions. However, in that work the condensate was represented through externally imposed, state-dependent potentials — an approximation that captures nonequilibrium transport but does not account for the back-reaction of chemistry on the phase-separated structure itself. Here, we develop a fully self-consistent formulation. We first recover the case in which a phase-separated scaffold controls the transport of chemically active client molecules, and we show that the same rules as before decide the presence of spatial fluxes, then extend the framework to active scaffold systems in which the phase-separating species themselves participate in the reaction network. This allows us to study how chemical activity reshapes condensate stability, modulates client partitioning, and generates directed transport across phase boundaries.

## I. MODEL DEFINITION

We consider a system of *N* particles, each of which can occupy one chemical state in the set

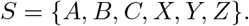

These states may represent distinct allosteric configurations of the same protein, or alternatively different chemical species. Since our focus is on a client–scaffold system, we interpret the states *A, B, C* as client states and the states *X, Y, Z* as scaffold states. Chemical transitions are allowed only between states belonging to the same physical role: for example, a particle in state A may transition to state B, but not to state *X*. We therefore define the total number of client and scaffold particles as *N*client and *N*scaffold, respectively, with

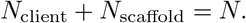

Consistently with other client–scaffold models, we focus on the dilute-client regime,

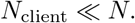

The distinction between client and scaffold states will become explicit through the choice of interaction and kinetic parameters, as described below.

We denote by *ρ*_*σ*_(*x*) the concentration field of particles in state *σ* ∈ *S*, and collect all concentration fields into the vector

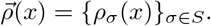

We consider a closed system, so that the total number of particles is conserved:

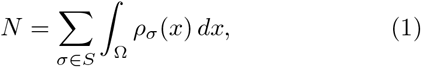

where Ω is the spatial domain. In the main text, we restrict the analysis to a one-dimensional domain,

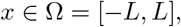

while in Appendix A we extend our results for a two-dimensional case.

The time evolution of the concentration fields is governed by a reaction–diffusion equation^27^,

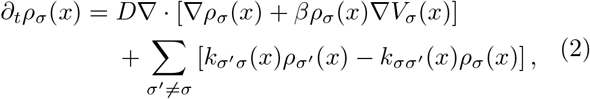

where *D* is the diffusion coefficient and *β* = (*kBT*) ^*−*1^ is the inverse thermal energy. The first term in Eq. (2) describes transport due to diffusion and drift, whereas the second term accounts for local transitions between chemical states. These two contributions are coupled through the conditional free energy *V*_*σ*_ (*x*): its gradient generates the drift force, while the reaction rates depend on the same local energetic landscape. In this framework, *V*_*σ*_ (*x*) represents the mean-field energetic contribution experienced by a particle in state *σ* at position *x* due to the remaining particles in the system^27^. A detailed derivation of Eq. (2), and of the role played by the conditional free energy, can be found in the original work by Shelest et al. ^27^.

In the following, we describe how the diffusion–drift and reaction terms are constructed self-consistently.

### A. Diffusion–drift term

The diffusion–drift term in Eq. (2) can be related to a generalized Cahn–Hilliard dynamics. Suppose that the system is described by a Helmholtz free-energy functional 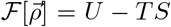, where U and *S* are the internal energy and entropy, respectively. The chemical potential of state *σ* is then defined as

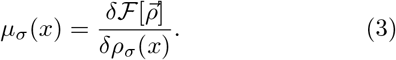

Assuming an ideal-gas entropic contribution, the chemical potential can be written as

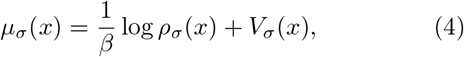

up to an additive constant. Here, the conditional free energy is the excess contribution to the chemical potential beyond the ideal entropic term:

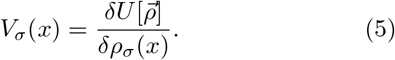

It follows that the diffusion–drift contribution to Eq. (2) can be written as

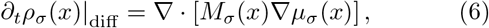

with density-dependent mobility

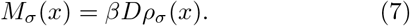

Thus, the transport part of Eq. (2) corresponds to a generalized Cahn–Hilliard equation with constant diffusion coefficient *D* and density-dependent mobility^22, 23^.

In Shelest et al.^27^, the conditional free energy was treated as an externally prescribed potential mimicking the effect of a condensate on client particles. Here, by deriving *V*_*σ*_ from a free-energy functional, we extend the formalism to describe both scaffold phase separation and client partitioning in a self-consistent manner.

We consider a free-energy functional of the form

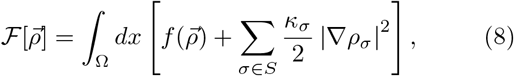

Where 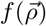 is the local free-energy density and *κ* _*σ*_ is the interfacial penalty associated with state *σ*. The gradient term penalizes sharp spatial variations in the concentration profiles and therefore sets the energetic cost of interfaces.

In this work, we use a Flory–Huggins free-energy density framework, which is the basis of many theoretical models of biological phase separation, while in Appendix B we show how the same formalism can be applied to the expansion of the free-energy around a critical point, leading to a Ginzburg–Landau description. Other free-energy models could also be incorporated, for example to describe more specific microscopic interactions.

The Flory–Huggins free-energy density used here is thus^21^

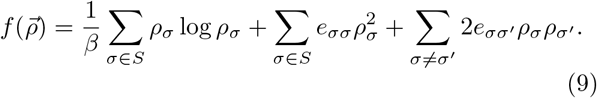

The first term is the ideal mixing contribution, while the remaining terms describe effective pairwise interactions between chemical states and define the local contribu-tion to the internal energy. The coefficients *e*_*σσ*_ and 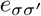 denote self- and cross-interaction strengths, respectively. Using Eq. (5), the conditional free energy of state *σ* is

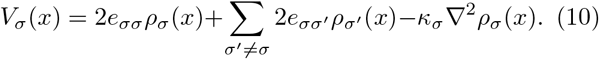

Thus, *V*_*σ*_ depends on the local interaction strengths, the concentration profiles, and the interfacial penalty through the Laplacian term.

In this work, we choose the interaction parameters according to the role played by each chemical state. Client states interact only with scaffold states, whereas scaffold states may interact both with themselves and with other scaffold states. This choice reflects the physical situation of interest: clients partition into or away from a scaffold-rich condensate, while the scaffold states provide the interactions required for phase separation.

### B. Reaction term

We now turn to the kinetic part of Eq. (2), which describes transitions between chemical states. Following Shelest et al. ^27^, we model these transitions as activated processes over kinetic barriers, in the spirit of Kramers barrier-crossing theory and transition-state theory^28, 29^. For a transition from state *σ* to state *σ* ^*′*^, we introduce an intermediate transition state *σσ* ^*′*^ and write the local reaction rate as

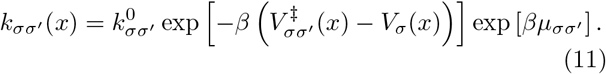

Here, 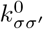 is a bare rate constant, 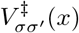 is the conditional free energy of the transition state, and 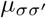 is the chemical potential supplied by the fuel. In the absence of fuel, 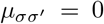 , the rates obey detailed balance because the forward and reverse transitions share the same transition-state barrier and bare rates^30, 31^. Nonzero values of 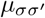 therefore provide a controlled mechanism for driving the chemical network out of equilibrium.

For the activated-process interpretation to remain valid, the transition state must lie above both reacting states. Therefore, the conditional free energy of the intermediate state must satisfy the conditions

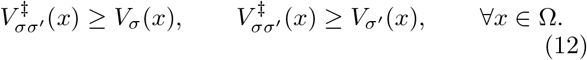

Whenever 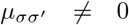, we treat the fuel chemical potential as spatially uniform. This corresponds to a chemostatted-fuel approximation commonly used in nonequilibrium thermodynamic descriptions of open chemical reaction networks^31^. In biological settings, the relevant fuel is often ATP or some other small molecule, such GTP or NADH, whose concentration is maintained by cellular metabolism and is typically much larger than the concentration of the molecular species undergoing the active process. A spatially uniform fuel chemical potential is therefore a natural first approximation: ATP is typically present at millimolar concentrations, it diffuses fast and is strongly buffered by cellular metabolism, so global ATP levels often remain nearly homeostatic despite large changes in turnover^32, 33^. Spatially heterogeneous fuel concentrations could be incorporated by treating nucleotides explicitly as additional particles in the system, with ATP and ADP represented as distinct chemical states. We leave this extension for future work and restrict ourselves here to a uniform chemostat.

In the simulations presented below, transitions are allowed only between states with the same physical role. States may interconvert within each species, but clients and scaffolds do not interconvert. Under this restriction, the free-energy functional determines the conditional free energies that enter the diffusion–drift dynamics, while the transition-state barriers remain an independent kinetic ingredient.

We therefore prescribe the barrier profiles phenomenologically, while enforcing the activated-process constraint above. For transitions between client states, *σ, σ* ^*′*^ ∈ {*A, B, C*}, we choose

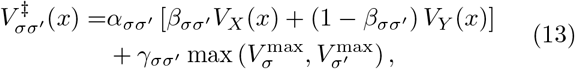

Where

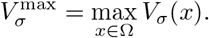

Here, 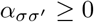 controls the amplitude of the spatially varying component of the barrier, 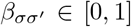 controls its shape by interpolating between two conditional free-energy profiles, and 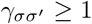 sets the constant offset required to keep the barrier above the reacting states. For client states, this choice models the fact that client interconversion reactions can be affected by the condensate environment^10, 34^. We therefore construct the client reaction barriers by interpolating between the spatial profiles of the scaffold states *X* and Y . In particular, 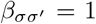 implies that the barrier follows the profile of *V*_*X*_ (*x*) and is therefore larger outside the condensate, where *V*_*X*_ (*x*) takes higher values. Conversely, 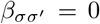 implies that the barrier follows the profile of *V*_*Y*_ (*x*) and is larger inside the condensate when phase separation occurs. Intermediate values of 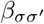 interpolate between these two limiting cases.

For transitions between scaffold states,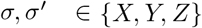, the barriers are modeled in an analogous way, but depend self-consistently on the conditional free-energy profiles of the reacting scaffold states:

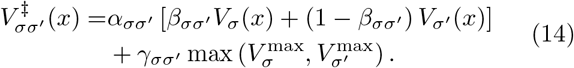

The model is therefore fully specified by the free-energy functional, the allowed chemical transitions, the fuel chemical potentials, and the transition-state barriers. We use this framework to investigate the interplay between phase separation, active chemical reactions, and diffusive fluxes. To this end, we solve Eq. (2) at steady state, meaning

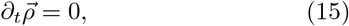

under different thermodynamic and kinetic conditions and for periodic boundary conditions. A full description of the numerical methods is provided in Appendix C, and reference parameters are shown in Table I unless explicitly stated.

**TABLE 1.** Model parameters used in the simulations. For parameters defined over multiple species or pairs, the ordering of the reported values is indicated in the table. Unless otherwise specified, the bare reaction rate constant for clients and scaffolds reactions are 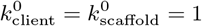

| Parameter group | Symbol | Order | Values |  |  |
| --- | --- | --- | --- | --- | --- |
| Transport and domain | $D, \beta, L$ | – | 1.0 | 1.0 | 100 |
| Scaffold self-interactions | $e_{\sigma\sigma}$ | $X, Y, Z$ | –0.01 | 0.10 | 0.05 |
| Scaffold cross-interactions | $e_{\sigma\sigma'}$ | $XY, XZ, YZ$ | 0.20 | 0.40 | –0.02 |
| Client–$X$ interactions | $e_{\sigma X}$ | $A, B, C$ | 0.001 | –0.02 | –0.02 |
| Client–$Y$ interactions | $e_{\sigma Y}$ | $A, B, C$ | –0.02 | 0.02 | 0.03 |
| Client gradient penalties | $\kappa_{\sigma}$ | $A, B, C$ | 0 | 0 | 0 |
| Scaffold gradient penalties | $\kappa_{\sigma}$ | $X, Y, Z$ | 0.5 | 0.5 | 0.5 |
| Client barrier amplitudes | $\alpha_{\sigma\sigma'}$ | $AB, AC, BC$ | 1.2 | 1.3 | 1.5 |
| Client barrier interpolation | $\beta_{\sigma\sigma'}$ | $AB, AC, BC$ | 0.5 | 0.5 | 0.5 |
| Client barrier offsets | $\gamma_{\sigma\sigma'}$ | $AB, AC, BC$ | 2.0 | 2.0 | 2.0 |
| Scaffold barrier amplitudes | $\alpha_{\sigma\sigma'}$ | $XY, YZ, ZX$ | 1.2 | 1.3 | 1.5 |
| Scaffold barrier interpolation | $\beta_{\sigma\sigma'}$ | $XY, YZ, ZX$ | 1.0 | 1.0 | 1.0 |
| Scaffold barrier offsets | $\gamma_{\sigma\sigma'}$ | $XY, YZ, ZX$ | 1.0 | 1.0 | 1.0 |

## II. RESULTS

We now use the self-consistent conditional free-energy framework to investigate how phase separation, chemical activity, and diffusive transport are coupled at steady state. Unless otherwise stated, the scaffold parameters are chosen such that the system forms a scaffold-rich condensate. We quantify phase separation through the density contrast of scaffold state *X*,

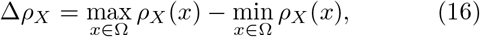

and quantify transport through the integrated absolute diffusive flux

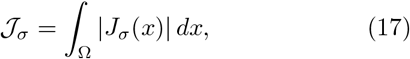

Where

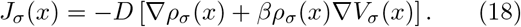

While 𝒥 _*σ*_ measures the total amount of transport, the spatial profiles of *J*_*σ*_ (*x*) retain information about the direction of transport across the condensate interface.

### A. Active client reactions generate interfacial fluxes

We first consider a regime in which the scaffold species form a stable condensate and fuel is applied only to the client reaction *A* →*B*, so that *µ* _*AB*_ ≠0. Figure 2a shows the resulting steady-state scaffold profiles. Because of its self-attraction, state *X* is strongly enriched in the central region of the domain, whereas the other scaffold states, *Y* and *Z*, which are repelled by *X* and attract each other (see Table I), are depleted from this region. We therefore identify the *X*-rich phase as the condensate.

**FIG. 1.**
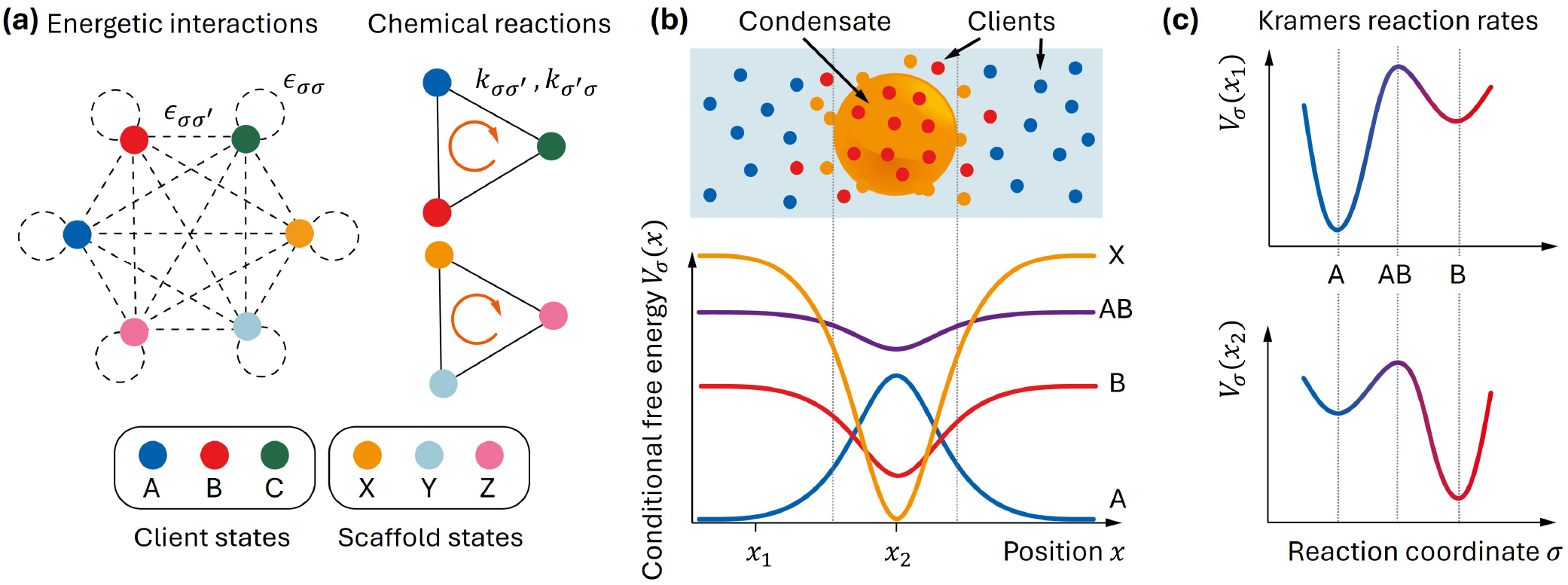
Schematic of the conditional-free-energy framework. (a) The system contains three client states, *A, B*, and *C*, and three scaffold states, *X, Y* , and *Z*. Self- and cross-interactions between states define the enthalpic contribution to the free energy, while chemical transitions are allowed only between states with the same role, either client or scaffold. (b) The conditional free energies derived from the free-energy functional determine both scaffold phase separation and client partitioning. In the schematic, scaffold state *X* forms a condensate, and the client states distribute according to their interactions with the scaffold-rich and dilute phases. (c) Chemical transitions are modeled as activated processes over state-dependent kinetic barriers. For a transition between states *A* and *B*, the intermediate state *AB* defines the transition-state barrier. Because both the reactant free energies and the barrier depend on position, the reaction rates are spatially heterogeneous.

**FIG. 2.**
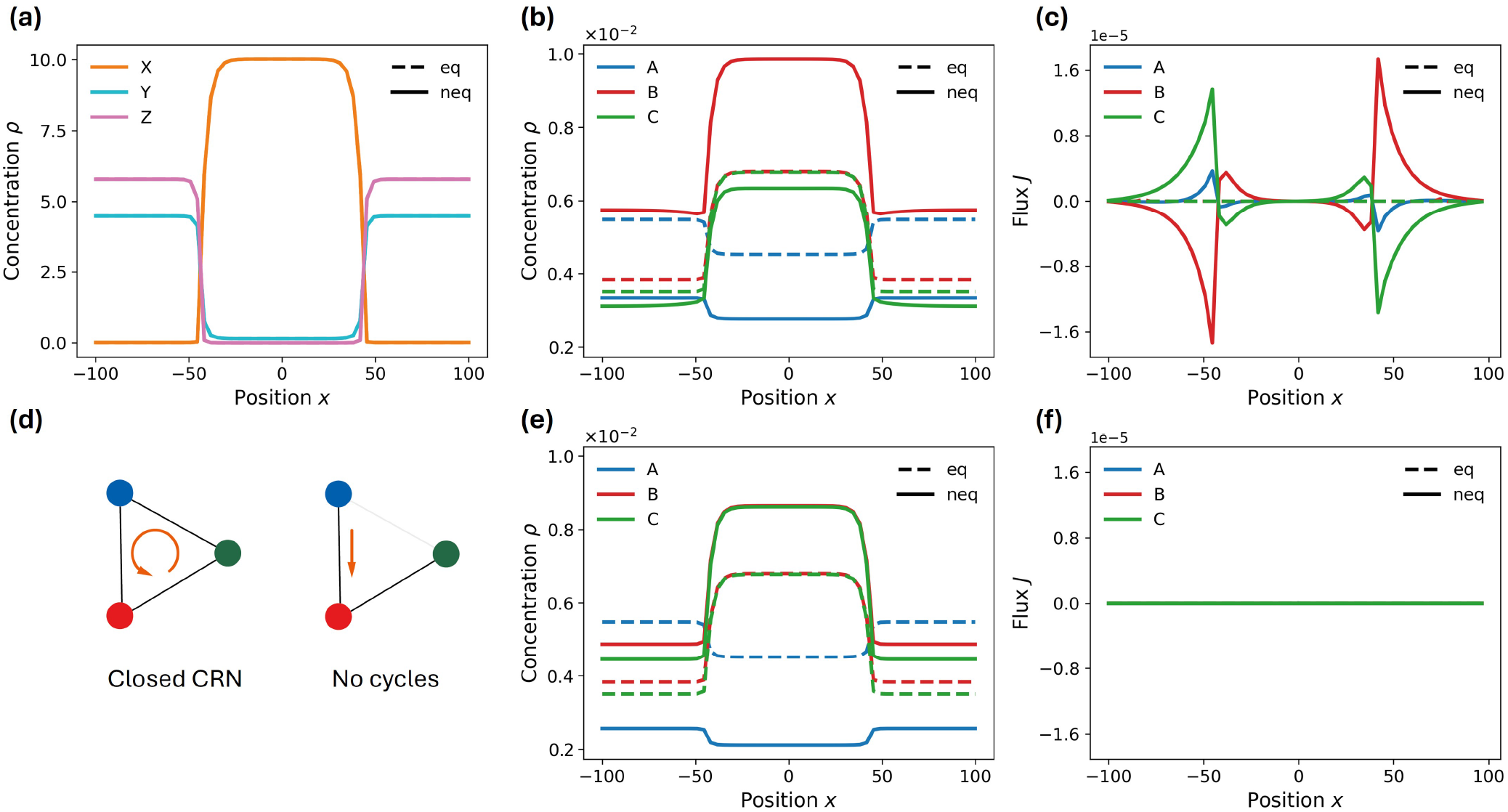
Active client chemical reaction cycles generate interfacial fluxes. (a) Steady-state concentration profiles of the scaffold states. State *X* forms a scaffold-rich condensate in the center of the domain, while states *Y* and *Z* are depleted from the condensate region. (b) Steady-state concentration profiles of the client states at equilibrium, dashed lines, and under nonequilibrium driving, solid lines. Driving the client reaction *A→ B* modifies client partitioning while leaving the scaffold condensate essentially unchanged. (c) Corresponding client diffusive fluxes. At equilibrium, detailed balance and boundary conditions imply vanishing fluxes. Under nonequilibrium driving, finite client fluxes emerge and localize at the condensate interfaces. (d) The client system is also studied for the case of an open chemical network by setting the bare reaction rate between states A and C to zero. The same fuel is applied to the reaction *A → B* as in the closed CRN case. (e) Steady-state concentration profiles for an open client network. (f) Corresponding client diffusive fluxes. Unlike the closed network case, here fluxes are always zero, independently of the fuel applied. Parameters are reported in Table I; here *N*_scaffold_ = 2000, *N*_client_ = 3 and *µ*_*AB*_ = 1.

The corresponding client profiles are shown in Fig. 2b. At equilibrium, clients partition between the dilute and condensed phases according to their interactions with the scaffold. For example, since *e*_*AX*_ > 0, state A is disfavored inside the *X*-rich condensate and preferentially localizes in the dilute phase. When the *A*→ *B* reaction is driven out of equilibrium, the scaffold condensate remains essentially unchanged because the clients are dilute, but the client profiles are modified.

The associated diffusive fluxes are shown in Fig. 2c. At equilibrium, detailed balance implies that the local reaction currents vanish (in agreement with previous results^27^ and trivially consistent with basic thermodynamics). The steady-state equation therefore reduces to

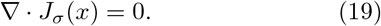

Together with periodic boundary conditions and consistently with basic thermodynamics, this gives *J*_*σ*_ (*x*) = 0 for every state. Equilibrium steady states therefore cannot sustain diffusive fluxes.

By contrast, under nonequilibrium driving, finite steady-state client fluxes emerge. These fluxes are localized primarily at the condensate interfaces, consistent with the mechanism previously identified for active reaction networks in spatially inhomogeneous systems^27^. Depending on the client state, the flux may point either into or out of the condensate, showing that active reactions can sustain directed exchange between the dilute and condensed phases.

We also investigate the role of the client network topology in determining the presence of NESS (nonequilibrium steady-state) fluxes. In particular, we repeat the simulation under the same parameter choice for an open client network as shown in Fig. 2(d). In order to do so, we simply set the bare rate constant between states A and C to zero, namely 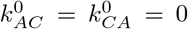, while still applying the fuel to the reaction branch *A* → *B*. Corresponding concentration profiles shown in Fig. 2(e) differ from the case of the closed client network (Fig. 2(b)). More importantly, they differ from the equilibrium case, when no fuel is applied to the *A* → *B* reaction. This is expected since the steady-state solution for the two systems is different. Nonetheless, this change due to the non-equilibrium driving of the open network does not result in NESS fluxes. This can be understood because a CRN (chemical reaction network) without cycles is solved using an effective detailed balance: the non-equilibrium driving only has the effect of changing some of the parameters of the system, which otherwise functions as if it were at equilibrium. This result is in agreement with the work of Shelest et al. ^27^.

### B. Kinetic barriers tune both the magnitude and direction of client transport

We next investigate how the kinetic barriers control nonequilibrium transport. In Fig. 3a, we vary the barrier-amplitude parameters α _*AB*_ and α _*AC*_, while constraining *α* _*BC*_ so that *α* _*BC*_ = *α* _*AC*_ and measure the integrated flux of species *B*,

**FIG. 3.**
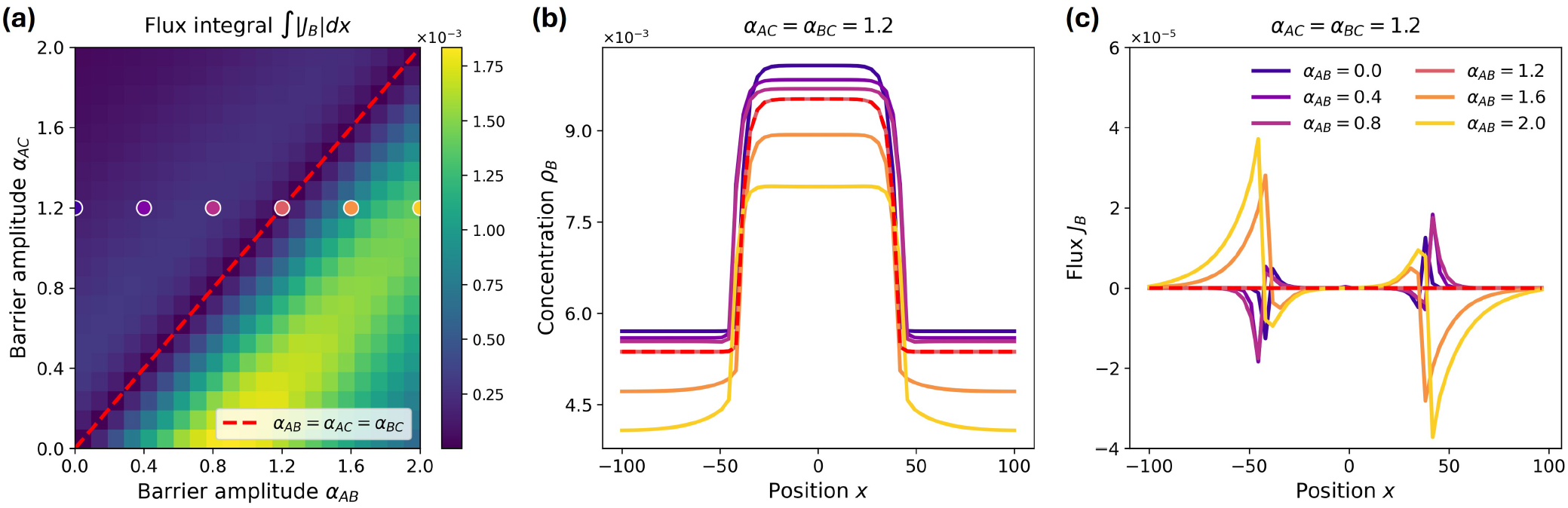
Kinetic barriers control the magnitude and direction of active client transport. (a) Integrated absolute flux of client state *B*, 𝒥_*B*_ = ∫_Ω_ |*J*_*B*_ (*x*) | *dx*, as a function of the client barrier amplitudes *α*_*AB*_ and *α*_*AC*_ , with *α*_*BC*_ fixed. The red dashed line marks the case *α*_*AB*_ = *α*_*AC*_ = *α*_*BC*_ , along which the flux is suppressed since all kinetic barriers share the same spatial dependence. (b) Steady-state concentration profiles of state *B* for selected values of *α*_*AB*_ , with *α*_*AC*_ = *α*_*BC*_ = 1.2 fixed. The dashed red curve indicates the *α*_*AB*_ = *α*_*AC*_ = *α*_*BC*_ case. (c) Corresponding flux profiles *J*_*B*_ (*x*). Changing only *α*_*AB*_ modifies both the magnitude and the sign of the interfacial flux, showing that the same driven reaction network can either import or export client state *B* depending on the kinetic barrier landscape. Parameters are reported in Table I; here *N*scaffold = 2000, *N*client = 3 and *µAB* = 1.

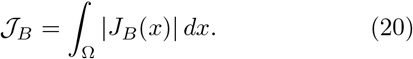

Similar, complementary results would be obtained for species *A* and *C*.

The heatmap shows that the flux depends strongly on the relative values of the barrier amplitudes. In particular, transport is suppressed along the diagonal *α* _*AB*_ = *α* _*AC*_, highlighted by the red dashed line. Along this diagonal, all the amplitude parameters for the kinetic barriers are the same, namely *α* _*AB*_ = *α* _*BC*_ = *α* _*AC*_. Since we also set the interpolation parameters to be the same, *β* _*AB*_ = *β* _*BC*_ = *β* _*AC*_ = 1/2, along the diagonal the conditional free energy of all barriers across clients reactions have the same spatial dependence, aside for a constant offset.

We have thus found that all the rules identified in Shelest et al. ^27^ are also valid in the present self-consistent scheme: the reaction network must contain a closed cycle (Fig. 2f), at least one transition must be driven out of equilibrium (Fig. 2c), and distinct reaction barriers must have different spatial dependence (Fig. 3a,c).

Figures 3b,c illustrate the effect of changing a single kinetic parameter. We fix α _*AC*_ = 1.2 and vary α _*AB*_. This modification is sufficient to change the steady-state profile of species B. More importantly, the corresponding flux profiles show that the sign of the interfacial current can reverse as α _*AB*_ is varied. Thus, for the same applied fuel and energies of the different interconverting states, the active reaction network can either drive species B into, or expel it from, the condensate, depending only on the kinetic barrier landscape.

These results highlight how imposing only detailed balance on the structure of the reactions, caring only for the free-energy difference between the interconverting states but disregarding the details of the reaction barriers, might miss crucial features of the systems under investigation, possibly leading to wrong results.

We have therefore established that a self-consistent description, in which the phase-separating scaffold provides the background experienced by the chemically active, non-phase-separating species, reproduces the results and obeys the same principles previously identified^27^. In particular, the externally imposed potential used in earlier treatments corresponds to the limit considered here in which scaffold particles greatly outnumber the client species. When the two populations become comparable, the energetic contribution arising from client– scaffold interactions becomes comparable to that arising from scaffold–scaffold interactions. As a result, clients can feed back on the scaffold and potentially induce non-trivial effects on phase behavior. These effects are beyond the scope of the present work and are left for future extensions.

### C. Active scaffold reactions shift the system across the phase-separation boundary

We then consider the case in which fuel is applied directly to a scaffold reaction. In this setting, activity acts on the species responsible for phase separation. As observed in the previous sections, at equilibrium the scaffold system phase separates into an X-rich condensate, while states Y and Z localize in the dilute phase outside of the condensate. Here, we apply the fuel to the reaction branch *X* → Y . This forces the system to a nonequilibrium steady-state where the state X is progressively depleted, ultimately affecting phase separation.

Figure 4a shows the density contrast Δ*ρ* _*X*_ as a function of the total particle number *N*_scaffold_ and the scaffold fuel chemical potential *µ* _*XY*_ . At low fuel and sufficiently large *N*_scaffold/_, the system remains phase separated. As *µ* _*XY*_ increases, however, the density contrast decreases, the condensate shrinks (Fig. 4b), and the system eventually becomes homogeneous. Fig. 4c suggests that this phenomenon is a first-order phase transition, as typical of phase separation away from the critical point, controlled by the fuel chemical potential *µ* _*XY*_ , an intensive quantity. We can then define a transition fuel value *µ* _*XY*_ (*N*_scaffold_), below which phase separation is preserved.

**FIG. 4.**
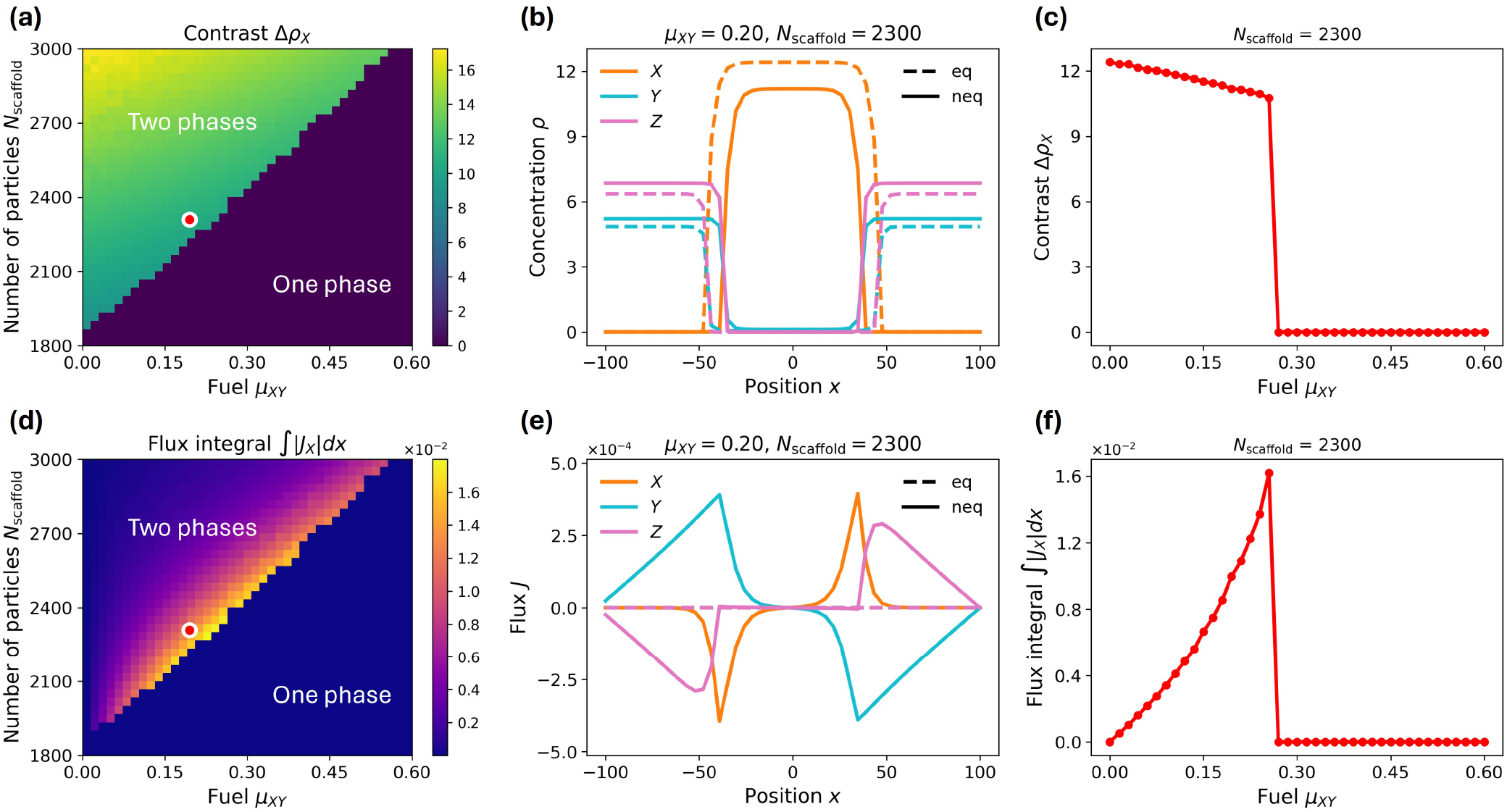
Active scaffold reactions remodel condensate stability and generate scaffold fluxes. (a) Density contrast of scaffold state *X*, Δ*ρ* _*X*_ , as a function of the total scaffold particle number *N*_scaffold_ and the fuel chemical potential applied to the scaffold reaction, *µ*_*XY*_ . Increasing scaffold activity shifts the system from a phase-separated regime to a homogeneous regime. (b) Representative scaffold concentration profiles at *N*_scaffold_ = 2300 and *µ* _*XY*_ = 0.2. Dashed lines show the corresponding equilibrium profiles, while solid lines show the nonequilibrium steady states. Increasing *µ*_*XY*_ weakens the *X*-rich condensate and reduces the density contrast. (d) Integral of the absolute value of scaffold *X* fluxes, 𝒥_*X*_ = ∫_Ω_ |*J*_*X*_ (*x*) | *dx*, over the same parameter space as in panel (a). The integral is maximal near the boundary between the phase-separated and homogeneous regimes. (e) Representative scaffold flux profiles corresponding to panels (b). At equilibrium, the fluxes vanish, whereas nonequilibrium scaffold driving generates finite fluxes localized around the condensate interfaces. (f) Integral of the absolute value of the diffusive flux plotted at different values of fuel *µXY* and for fixed *N*scaffold = 2300. Parameters are reported in Table I.

Analogously, we can analyze the integral of the diffusive flux of state X, 𝒥 _*X*_ (Fig. 4d). 𝒥*X* is zero in the homogeneous regime, as expected since there are no spatial variations of the concentrations. It also trivially vanishes when *µ* _*XY*_ vanishes, since the system is at equilibrium. More interestingly, it reaches the largest values at the boundary (Fig. 4e). This is particularly evident in Fig. 4f, where it is possible to see an increasing trend in the integrated flux up to the transition, above which the flux drops to zero discontinuously. This intriguing result is due to the dual role of the non-equilibrium drive: on the one hand, the larger the drive (*µ* _*XY*_), the more energy to support fluxes; on the other hand, the first-order phase transition is such that the switch to the homogeneous phase is abrupt: 𝒥 _*X*_ can thus keep increasing up to the transition. In the presence of a second-order phase transition it could have been expected a more complex behavior of the flux, first increasing with the fuel, then decreasing following the gradual change toward homogeneous behavior, but the exploration of this behavior goes beyond the scope of the present work.

These results suggest that away from the critical point, where the transition becomes continuous, the edge of condensate stability is a particularly efficient regime for sustaining active transport, consistent with the emerging view that interfacial properties can dominate nonequilibrium condensate behavior^26, 27^.

### D. Client and scaffold reaction cycles are coupled through the condensate environment

Finally, we drive both client and scaffold reactions out of equilibrium. Figure 5 summarizes the resulting steady states as a function of the client fuel µ*AB* and the scaffold fuel *µ* _*XY*_ .

**FIG. 5.**
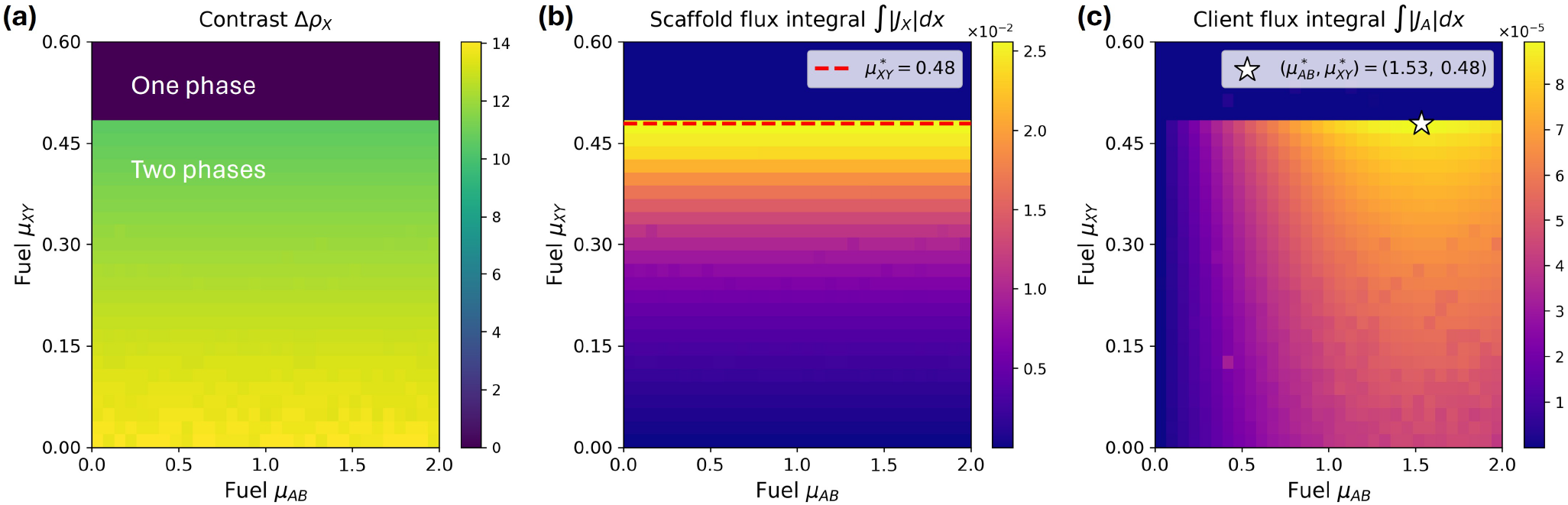
Client and scaffold reaction cycles are coupled through the condensate environment. (a) Density contrast of scaffold state *X*, Δ*ρ*_*X*_ , as a function of the fuel chemical potentials applied to the client reaction, *µ*_*AB*_ , and to the scaffold reaction, *µ*_*XY*_ . The condensate stability is controlled primarily by scaffold driving. (b) Integral of the absolute value of scaffold *X* flux, *J*_*X*_ = Ω |*J*_*X*_ (*x*)| *dx*, over the same parameter space. The scaffold flux is mainly controlled by *µ*_*XY*_ and is maxi mal near the loss of phase separation, indicated by the dashed red line. (c) Integral of the absolute value of client *A* flux, 𝒥 _*A*_ = Ω |*J*_*A*_(*x*)| *dx*.Client transport increases with direct client driving but is also enhanced near the scaffold-driven phase boundary. The star marks the maximum client flux at (*µAB, µXY*) = (1.53, 0.48). Parameters are reported in Table I. *N*_scaffold_ = 2500, *N*_client_ = 3.

The density contrast of the scaffold state *X* depends primarily on *µ* _*XY*_ and is only weakly affected by *µ* _*AB*_. This is expected because the scaffold species, whose number is much larger than that of the client species, determine the phase behavior, whereas the dilute client species do not substantially remodel the condensate in the parameter regime considered here.

The scaffold flux integral 𝒥 _*X*_ shows a similar dependence. It is controlled mainly by *µ* _*XY*_ and is maximal close to the loss of phase separation, consistent with the behavior observed when only the scaffold reaction is driven.

By contrast, the client flux integral 𝒥 _*A*_ depends on both fuel parameters. Increasing the client fuel enhances client transport, but the magnitude of this transport is also modulated by scaffold activity. In particular, 𝒥 _*A*_ is largest near the region where the scaffold condensate approaches the transition from a phase-separated to a homogeneous state. This indicates that scaffold activity indirectly controls client transport by reshaping the condensate environment.

Together, these results show that distinct active reaction cycles can be coupled through the shared spatial organization of the system. Scaffold activity controls the stability of the condensate, while client activity controls the conversion and redistribution of client states. Because client fluxes are enhanced near the phase-separation boundary, active remodeling of the scaffold condensate can regulate molecular-client transport without requiring direct changes to client-scaffold interactions.

Overall, the model reveals three central effects. First, active client reactions generate fluxes that localize at condensate interfaces. Second, the magnitude and direction of these fluxes are controlled by the kinetic barrier landscape. Third, active scaffold reactions can tune the system toward or away from phase separation, with maximal transport occurring near the edge of condensate stability. These results highlight the importance of treating thermodynamics and kinetics self-consistently when modeling chemically active condensates.

### E. Two-dimensional extension of the framework

In the preceding sections we have analyzed in some detail a one-dimensional setting for Eq. 2. Its extension to higher dimensions is straightforward, although the numerical integration is more cumbersome and time-consuming, and requires some further care. In Appendix A we show the results in two dimensions for a few representative cases, showing that the same phenomenology uncovered in one dimension remains unchanged. We expect differences in cases where the surface-to-volume ratio matters, possibly by choosing different condensate shapes.

## III. DISCUSSION AND CONCLUSION

Biomolecular condensates are often characterized through their equilibrium partitioning properties, namely their ability to enrich or exclude molecular species. This description is necessary, but incomplete for chemically active condensates. When reactions continuously inter-convert molecular states, phase separation, diffusion, and chemistry jointly determine the steady state, which may contain persistent fluxes across phase boundaries rather than static partition coefficients alone.

The central result of this work is that active condensates regulate molecular distributions and transport through two coupled forms of control. State-dependent interactions set the thermodynamic bias for localization inside or outside the condensate, whereas state-dependent transition barriers determine where interconversion preferentially occurs. At equilibrium, these biases fix molecular partitioning. Away from equilibrium, chemical activity can modify both partitioning and transport, with the resulting steady state depending on the spatial structure of reaction barriers as well as on free-energy differences between phases. Systems with similar equilibrium partitioning can therefore display distinct nonequilibrium profiles, including fluxes whose direction is only controlled by the kinetic barriers. In the regimes studied here, these fluxes localize near the condensate interface, identifying the boundary between dense and dilute phases as a privileged region for chemically driven transport.

When the scaffold itself participates in the reaction network, activity also feeds back on the condensate. Reactions no longer move clients through a fixed phase-separated background; they alter the stability of the phase-separated state. Increasing scaffold driving can weaken and eventually dissolve the condensate, while transport is maximal near the boundary between the phase-separated and homogeneous regimes. Condensates close to their stability threshold may therefore be especially sensitive to chemical regulation.

The framework developed here is deliberately minimal. We considered a one-dimensional system, a small number of client and scaffold states, and a spatially uniform fuel chemical potential. We also prescribed transition-state barriers phenomenologically, rather than deriving them from microscopic reaction coordinates. These simplifications allowed us to isolate the basic physical mechanisms, but they also point to natural extensions. Future work could include, for instance, higher-dimensional settings to explore surface-to-volume ratio effects or shape instabilities, explicit ATP/ADP dynamics, more complex reaction networks, and barrier profiles derived from microscopic models or inferred from experiments.

Despite these limitations, the model provides a general route for studying active condensates beyond equilibrium partitioning. It shows how chemical activity can generate interfacial fluxes, tune the stability of phase-separated compartments, and couple distinct reaction cy-cles through a shared condensate environment. This may be relevant for biological condensates that function not only as selective compartments, but also as active processing centers that organize molecular flows in space.

## AUTHOR CONTRIBUTIONS

Both authors conceived the research, analyzed the results and wrote the manuscript. MDM wrote the codes and performed the numerical calculations.

## DATA AVAILABILITY

The data that support the findings of this study are available within the article.

## Appendix A Two-dimensional extension

We replicate the same results presented in the main text for a two-dimensional setting. In this section we show that the results regarding the active regulation of phase behavior and that maximal diffusive fluxes localize in the phase space at the stability edge of the condensate also apply to two-dimensional systems, thus can be interpreted as general features of the model.

Fig. 6 shows the results for the two-dimensional system analogous to the active-scaffold case studied in Section II C. The system is driven out of equilibrium by applying fuel *µXZ* to the reaction *X*→*Z*. As in the one-dimensional case, the applied fuel has a strong effect on phase behavior (Fig. 6a). Large fuel values shrink the condensate (Fig. 6e,f) and eventually suppress phase separation, in agreement with the one-dimensional results. Here, however, we also observe an additional effect. For values of *N*scaffold that do not support phase separation at equilibrium, intermediate fuel values can enrich the condensate-forming state *X* sufficiently to induce phase separation (Fig. 6c,d). This suggests that, in some parameter regimes, condensate formation may require nonequilibrium conditions. This fuel-induced phase separation is likely related to the specific interaction parameters chosen for this numerical experiment, rather than being a generic feature of the two-dimensional model. A systematic exploration of the parameter space would be required to determine which emergent behaviors are robust in two dimensions; this analysis is beyond the scope of the present work.

**FIG. 6.**
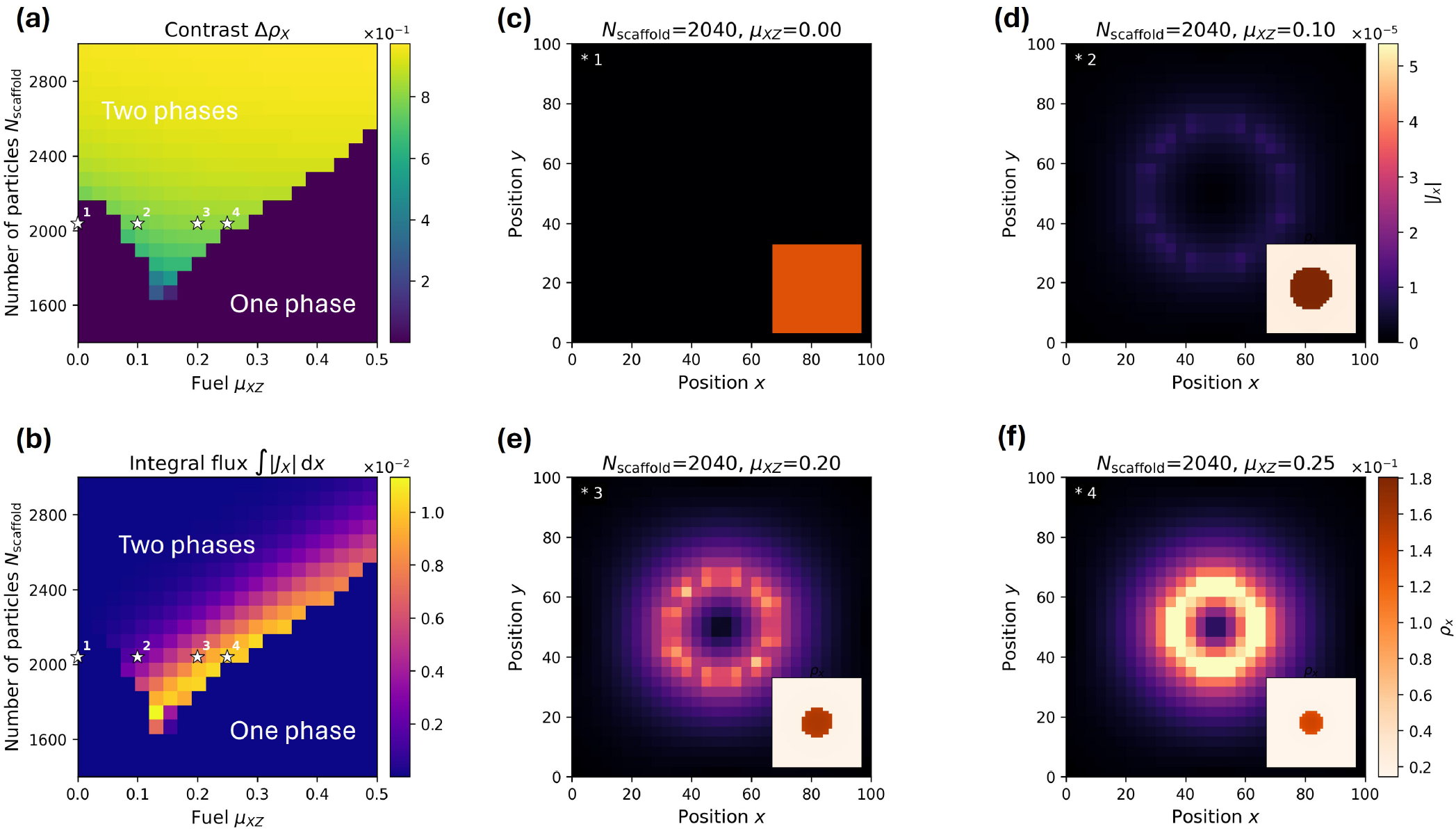
Active scaffold molecules in 2D. (a) Contrast for concentration of state X in the *N*scaffold vs fuel *µ* _*XZ*_ phase space. High-contrast regions represent the two-phase regime while zero contrast represents the homogeneous case. (b) Integral of diffusive fluxes of particle *X* in the two-dimensional domain. The fluxes are maximal in correspondence to the border between the phase separated and homogeneous regions. (c, d, e, f) Example concentrations (insets) and fluxes for fixed *N*_scaffold_ = 2040 and varying fuel values. In this case, phase separation arises only for intermediate values of fuel applied. Fluxes increase as we increase the fuel while still being in the phase separated region. Parameters used for this simulation are: *e*_*XX*_ = 0.5, *e*_*Y Y*_ = 1, *e*_*ZZ*_ = 8, *e*_*XY*_ = 10, *e*_*XZ*_ = 2, *e*_*Y Z*_ = 1, *e*_*AX*_ = 0.1, *e*_*BX*_ = *−* 2, *e*_*CX*_ = *−* 2, *e*_*AY*_ = *−* 2, *e*_*BY*_ = 2, *e*_*CY*_ = 3, *κ*_*X*_ = 0.1, *β*_*AB*_ = *β*_*AC*_ = *β*_*BC*_ = 1. All other parameters are as in Table I.

Finally, the behavior of the integrated absolute flux is also preserved in two dimensions: its maximal values remain localized in the phase diagram near the boundary between the homogeneous and phase-separated regimes (Fig. 6b).

## Appendix B Ginzburg–Landau free-energy description

The conditional free-energy framework introduced in the main text is not restricted to the Flory–Huggins form of the free-energy density; it applies to any functional for which the ideal entropic contribution can be separated from the excess. The Ginzburg–Landau description^35^ is a common choice in the biological phase-separation literature^24, 25^, as it is obtained by expanding the Flory–Huggins free energy around a critical concentration and often admits simple analytical solutions for the steady-state profiles. Here we show how to incorporate it into the present formalism and derive the corresponding conditional free energy explicitly. Writ-ing 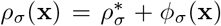 with 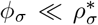 , and expanding the Flory–Huggins free-energy density, Eq. (9), to fourth order in *ϕ* _*σ*_ , one obtains

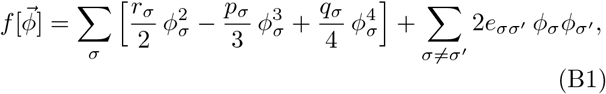

where the linear terms vanish by definition of 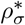, and the coefficients are given by

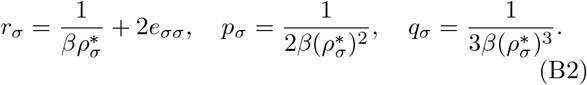

The cubic term in Eq. (B1) is generically present and vanishes only under additional symmetry conditions on 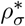 .

Within this description, the ideal entropic contribution 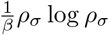 log is absorbed into the polynomial coefficients, so the conditional free energy cannot be isolated by direct functional differentiation of Eq. (B1). To recover it, we add and subtract the ideal entropic term inside the functional before differentiating, defining

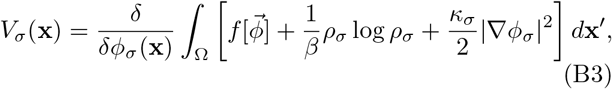

which yields

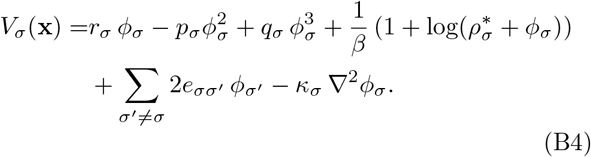

This expression can be substituted directly into Eq. (2), enabling a consistent treatment of transport and chemical transitions within the Ginzburg–Landau framework.

## Appendix C Numerical methods

We solve Eq. (2) directly at steady state using a Gummel-type fixed-point iteration scheme for nonlinear drift–diffusion systems,^36, 37^. Because the conditional free energies and reaction rates depend nonlinearly on the concentration profiles, at each iteration we freeze these quantities using the profiles from the previous step, solve the resulting linear system, and update the concentrations until convergence.

We discretize the one-dimensional domain Ω = [ − *L, L*] on *n*_*x*_ uniformly spaced grid points with spacing Δ*x*. The concentration fields are collected into a single vector

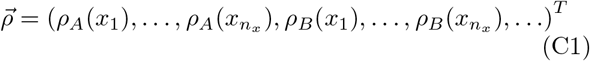

of size *n*_*x*_*n*_*σ*_ , where *n*_*σ*_ is the number of chemical states. At iteration *i*, the steady-state equation takes the linearized form

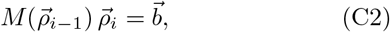

where *M* = *M* ^dd^ + *M* ^react^ is the sum of the diffusion– drift and reaction operators, assembled from the conditional free energies, transition-state barriers, and reaction rates evaluated at 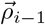. The diffusion–drift operator is discretized using a Scharfetter–Gummel scheme^37, 38^, which preserves the balance between diffusive and drift currents and remains stable when the drift term is large. Periodic boundary conditions are imposed on all concentration fields.

The right-hand side 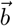 is zero except for the rows encoding mass-conservation constraints, which are required because the steady-state operator is otherwise singular. The form of these constraints depends on the topology of the reaction network. If no reactions are present, the particle number in each state is independently conserved, and we impose one constraint per species,

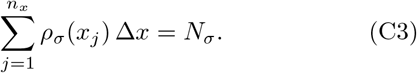

If all states are connected by reactions, only the total particle number is conserved, and a single global constraint suffices,

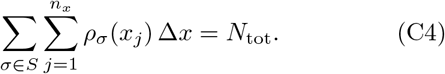

In the general case of a network with several disconnected components, one constraint is imposed per component. For the client–scaffold systems considered in the main text, this yields two constraints: one for the total number of client particles and one for the total number of scaffold particles.

Because the nonlinear fixed-point iteration can be stiff, we stabilize convergence using matrix mixing^39^. Rather than assembling the operator from the current iterate alone, we define the mixed operator

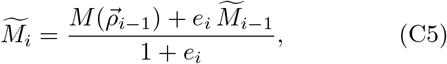

where *ei* ≥ 0 is a mixing parameter, and solve

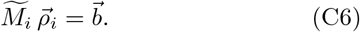

This damps rapid changes in the discretized operator between successive iterations and improves stability of the nonlinear solve. The iteration is continued until the relative change in the concentration vector falls below a prescribed tolerance,

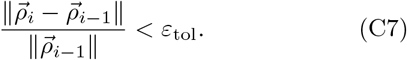

